# A Structure-Based hERG Pose Selection Workflow Integrating Induced-Fit Docking, Shape Similarity, Interaction Fingerprints, and Molecular Dynamics

**DOI:** 10.64898/2026.10.07.757389

**Authors:** Purvi Neema, Umut Ozuguzel, Saeed Najafian, Bodhisattwa Chaudhuri

## Abstract

The recent availability of several ligand-bound hERG/Kv11.1 cryo-EM structures creates an opportunity for prospective structure-based modeling, but it also introduces a practical problem: a new ligand may be compatible with only a subset of the available receptor conformations, and docking may generate several plausible poses within each template. We therefore evaluated a template-aware pose-selection framework using the astemizole-, pimozide-, and E4031-bound hERG structures as experimental reference environments, with apo 8ZYN included as a receptor-state control. The ligand set separated into two study-defined chemical regions: a low-polarity, highly lipophilic Group I represented by astemizole and pimozide, and a more polar Group II represented by E4031. Induced-fit docking (IFD) improved self-docking relative to rigid Glide, whereas apo 8ZYN did not reproduce ligand-bound behavior during short molecular dynamics (MD) assessment. Across the retrospective ligand-bound analyses, six of seven group-matched systems met the dual MD-quality criterion, whereas none of eight group-mismatched systems did. Highest static shape similarity was the most consistent pose-selection criterion for Group I, while an interaction-fingerprint (IFP)-favored 30:70 shape:IFP composite was retained provisionally for the single retrospective Group II reference. Application to dofetilide, ibutilide, and terfenadine produced binding hypotheses with residue-contact patterns consistent with independent hERG literature. Together, these results support using ligand-bound hERG structures as chemically selective reference environments for pose prioritization rather than as interchangeable receptor templates.

## 1. Introduction

The *KCNH2* gene encodes the human ether-à-go-go-related gene potassium channel (hERG/Kv11.1), which conducts the rapid component of the delayed rectifier potassium current (*I_Kr_*) and plays an important role in cardiac repolarization. Pharmacological inhibition of hERG can reduce *I_Kr_*, prolong ventricular repolarization and the QT interval, and increase the risk of Torsades de Pointes and other potentially life-threatening ventricular arrhythmias [1, 2]. Consequently, hERG liability is an important consideration during drug development. Quantitative structure-activity relationship, pharmacophore, and machine-learning models based on ligand information are widely used for early liability prediction [3, 4]. These approaches generally do not provide a protein-bound pose or explain how an individual compound interacts with specific residues in the channel. Structural modeling complements them by providing mechanistic hypotheses for ligand binding.

Structural modeling of hERG is challenging because chemically diverse blockers bind within a large central cavity formed by four equivalent subunits. Tyr652 and Phe656 on the S6 helices are major determinants of block for many inhibitors. Residues near the pore helix and the base of the selectivity filter, including Thr623, Ser624, and Val625, also contribute, depending on the ligand; Gly648, Ser649, and other inner cavity residues can be involved for some ligand classes [5–7] . Different chemotypes can therefore occupy the same general cavity while making overlapping but distinct residue interactions. The fourfold symmetry of the pore adds another source of ambiguity because symmetry-equivalent orientations may be biologically comparable even when they appear different in conventional coordinate comparisons [8] .

For many years, hERG binding modes had to be modeled without experimental structures containing bound inhibitors. Homology models, multiple receptor conformations, molecular dynamics, and rescoring after docking were used to explore plausible poses [8]. The structural basis has since improved substantially. The first full-length cryo-EM structure of open hERG established the overall architecture and central cavity geometry [9] , and a later structure with astemizole provided an early view of inhibitor binding [10] . More recently, Miyashita and co-workers resolved hERG structures bound to astemizole, pimozide, and E4031 together with an inhibitor-free apo structure (PDB 8ZYO, 8ZYQ, 8ZYP, and 8ZYN; [11]). These structures provide experimentally supported ligand poses and clearer local organization of residues in the pore helix, selectivity filter base, and S6 cavity, including Thr623, Ser624, Val625, Gly648, Ser649, Tyr652, and Phe656. They also provide receptor environments stabilized by different ligands that can be evaluated for prospective docking.

Having several experimental hERG structures creates a new question for prospective docking: which receptor structure is most appropriate for a ligand that does not have its own hERG complex? Sun, Wang, and Shen reported that the apo and ligand-bound structures are highly similar overall, with pairwise Cα RMSDs below 1 Å, while the Y652 side chains show substantial conformational flexibility within the binding pocket [12] . Local differences can therefore be small at the level of the whole protein but still matter within the cavity. A new blocker raises two practical questions: which ligand-bound receptor environment should be used as the starting structure, and which pose should be selected from the alternatives generated in that environment?

Docking score alone does not resolve the second question. Benchmark studies have shown that poses close to the experimental geometry can be generated without being ranked first by the scoring function [13] . Receptor flexibility also affects pose generation. Rigid Glide retains the deposited receptor geometry, whereas IFD allows local side-chain refinement during pose generation and redocking [14, 15] . Once a pose ensemble has been generated, experimentally resolved complexes provide additional information for ranking. Three-dimensional shape similarity measures how closely a candidate occupies the spatial volume of a reference ligand [16] , and interaction fingerprints compare residue interaction patterns [17]. Short MD simulations in a membrane environment can then examine whether a selected pose remains coherent after relaxation of the protein, ligand, membrane, solvent, and ions.

These considerations motivated the working framework examined in this study (Fig. 1). The available ligand-bound structures were treated as candidate receptor environments rather than automatically interchangeable templates. Physicochemical descriptors and key substructure features were used to characterize the ligands, IFD was used to generate candidate poses, and docking score, shape similarity, interaction fingerprint similarity, and composite shape:IFP criteria were compared for pose ranking before MD assessment. Within the chemical space represented by the current structures, Group I contains low-polarity, highly lipophilic ligands associated with the astemizole- and pimozide-bound environments, whereas Group II contains more polar ligands associated with the E4031-bound environment. The retrospective docking results were analyzed using this organization to examine template transferability and pose-selection behavior. The resulting strategy was then applied prospectively to dofetilide, ibutilide, and terfenadine. The objective was to determine whether ligand chemistry, receptor environment, reference-informed pose ranking, and dynamic assessment can be combined into a practical strategy for hERG pose prioritization when the modeled ligand lacks its own experimental hERG structure.

**Fig. 1.**
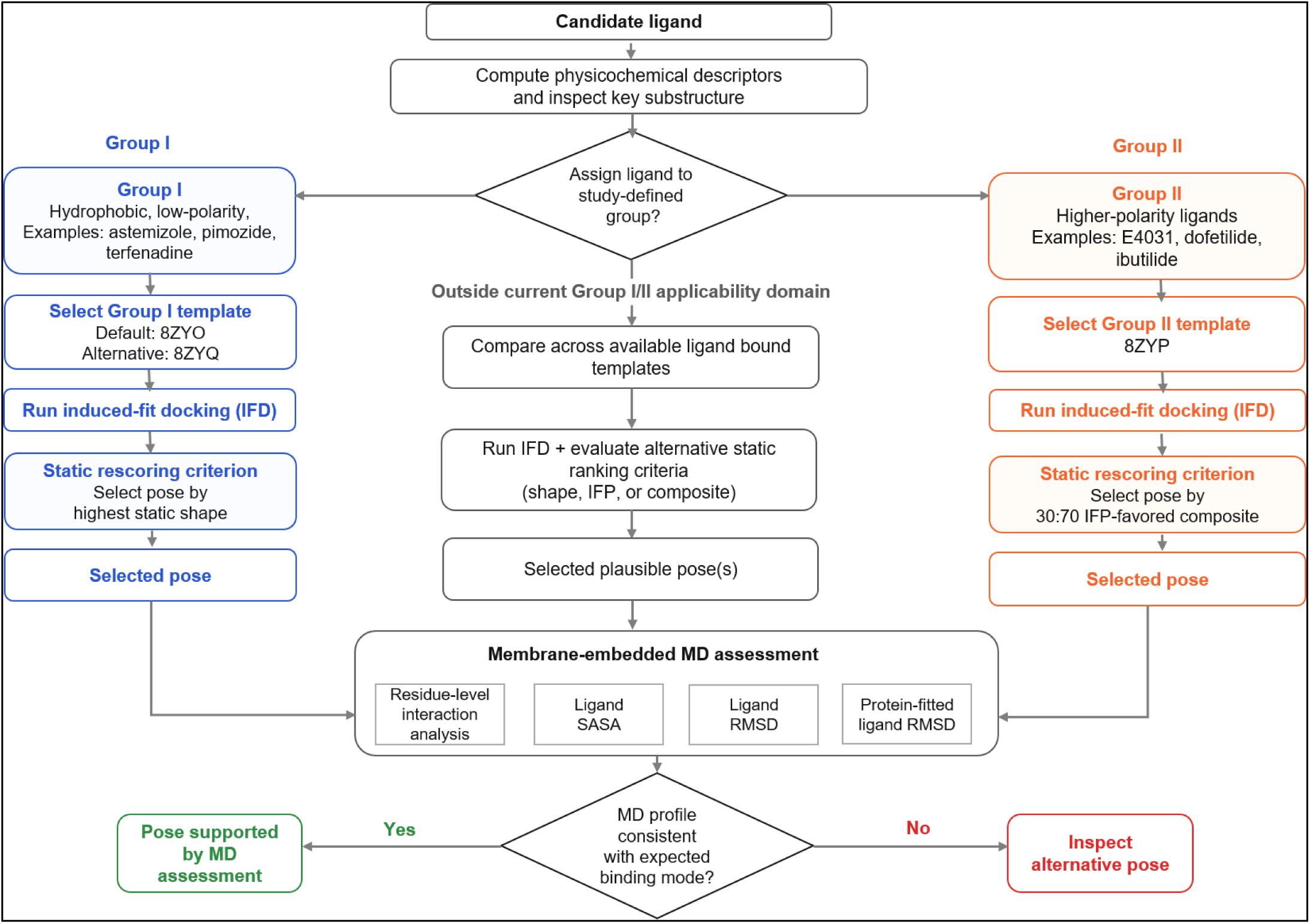
Proposed prospective framework for receptor-template selection, pose ranking, and MD assessment in hERG

## 2. Methods

### 2.1 Computational platform and overall workflow

All receptor and ligand preparation, docking, membrane system construction, molecular dynamics (MD) simulations, and trajectory analyses were performed in the Schrödinger Suite 2025-2 on the University of Connecticut high performance computing cluster. The OPLS4 force field was used for molecular preparation, docking refinement, and MD parameterization [18].

The computational implementation separated generation of candidate poses from pose selection and dynamic assessment. Docking generated alternative hERG inhibitor poses in the cryo-EM receptor templates. Six predefined pose-selection criteria were then applied: docking score, 3D shape similarity, static interaction fingerprint (IFP) similarity, and three shape:IFP composite scores. For the three reference inhibitors, the corresponding cryo-EM structures provided experimental reference poses for retrospective assessment. The highest ranked pose from each criterion was evaluated by short MD simulations in a membrane environment using interaction recovery, ligand burial, ligand RMSD, and protein-fitted ligand RMSD as descriptors after MD. The computational implementation is summarized in Supplementary Fig. S1.

### 2.2 Receptor structures and ligand preparation

Four hERG structures from the higher-resolution cryo-EM study of Miyashita and co-workers were used as receptor templates: astemizole-bound 8ZYO, pimozide-bound 8ZYQ, E4031-bound 8ZYP, and inhibitor-free apo 8ZYN [11]. For each ligand-bound structure, the resolved inhibitor was removed before docking. Receptors were prepared in Maestro using the Protein Preparation Workflow at pH 7.4, including hydrogen addition, hydrogen-bond network optimization, PROPKA-assisted protonation state assignment [19], and correction of overlapping hydrogens. The prepared receptors were then used for grid generation and induced-fit docking.

Astemizole, pimozide, E4031, dofetilide, ibutilide, and terfenadine structures were retrieved from PubChem [20] and prepared with LigPrep. LigPrep used the OPLS4 force field, and possible ionization and tautomeric states were enumerated using Epik Classic at pH 7.4 ± 2.0 [21]. TPSA and XLogP3-AA values used for ligand characterization were obtained from PubChem. The resulting prepared ligand sets were used as input to both rigid Glide docking and IFD.

### 2.3 Rigid Glide docking

Rigid receptor docking was performed with Glide in Standard Precision (SP) mode [14, 22]. Receptor grids were centered on the canonical hERG inner cavity binding region defined by Tyr652 and Phe656, with default inner and outer box dimensions used throughout. The receptor remained rigid during docking, and no positional, hydrogen bond, or pharmacophore constraints were applied.

Rigid Glide served as the comparator in the self-docking benchmark. Each inhibitor resolved by cryo-EM was docked back into its own receptor template: astemizole into emptied 8ZYO, pimozide into emptied 8ZYQ, and E4031 into emptied 8ZYP. The resulting Glide poses were compared with IFD poses after MD assessment.

### 2.4 Induced-fit docking

Induced-fit docking (IFD) was performed using the Schrödinger IFD workflow with the standard protocol, standard accuracy, and no additional constraints [15] . IFD allowed local receptor side-chain relaxation through Prime optimization followed by Glide redocking. The resulting IFD pose pools were used for self-docking, apo docking, ligand-referenced cross-docking, template-referenced cross-docking, and prospective docking of dofetilide, ibutilide, and terfenadine. The pose with the most favorable IFD score in each pool was retained as one of the predefined selection candidates.

### 2.5 Docking system design

Five docking scenarios were evaluated to separate pose generation, receptor-template dependence, and the effect of reference availability. (i) Self-docking: astemizole-8ZYO, pimozide-8ZYQ, and E4031-8ZYP, in which each ligand was docked into its cognate experimental receptor template and used for the Glide versus IFD comparison. (ii) Apo docking: astemizole, pimozide, and E4031 were docked into apo 8ZYN as a receptor state control. (iii) Ligand-referenced cross-docking: each reference ligand was docked into the other two ligand-bound templates and evaluated against its own cryo-EM reference pose. (iv) Template-referenced cross-docking: reference ligands were docked into the other ligand-bound templates, with the template-bound ligand and its interaction fingerprint used as the static reference. This represents the practical case in which the docked ligand has no cognate hERG cryo-EM pose. (v) Prospective applications: dofetilide, ibutilide, and terfenadine were each docked into 8ZYO, 8ZYQ, and 8ZYP. These scenarios are summarized in Table 1.

**Table 1.**
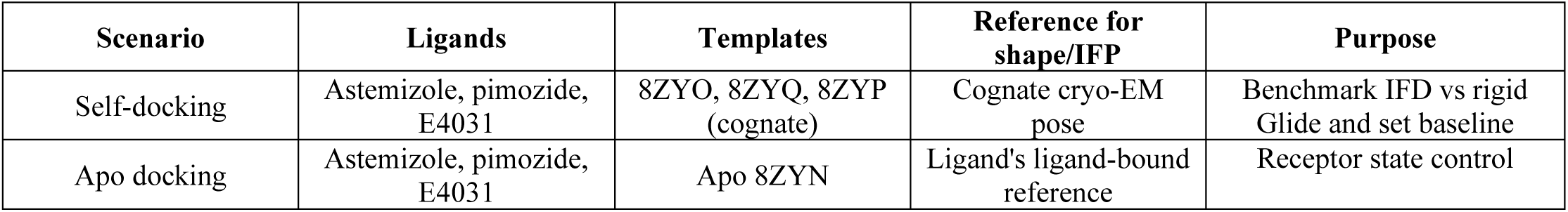
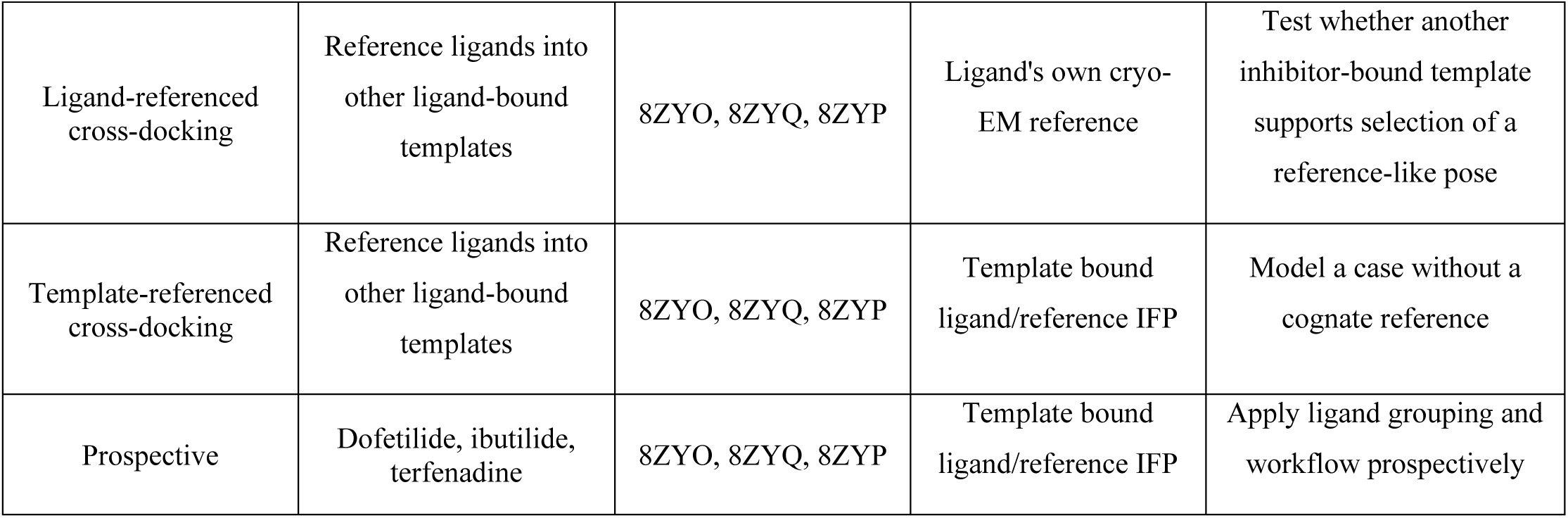
Summary of the docking scenarios evaluated in this study, including ligands, hERG receptor templates, structural references used for shape and interaction fingerprint comparisons, and the purpose of each docking experiment.

The reference used for shape and interaction fingerprint comparison depended on the docking scenario. In ligand-referenced cross-docking, each docked ligand was compared with its own experimentally resolved cryo-EM pose. In template-referenced cross-docking, the bound ligand and interaction fingerprint of the receptor template served as the reference. For apo docking, the shape reference was the docked ligand’s own cryo-EM pose, and the static IFP reference was its ligand-bound complex. Dofetilide, ibutilide, and terfenadine were treated as prospective ligands and compared with the bound ligand of each template.

### 2.6 Static 3D shape similarity ranking

Three-dimensional ligand shape similarity was calculated using ROSHAMBO2, which uses GPU acceleration for Gaussian shape overlap [23]. For each docking system, the reference ligand specific to that scenario was used as the query and docked candidate poses formed the comparison set. Each candidate was rigidly superposed with the reference by optimizing shape overlap, and similarity was quantified using the resulting shape Tanimoto coefficient. This comparison measures three-dimensional volumetric similarity without requiring identical atom-to-atom correspondence. The candidate with the largest shape Tanimoto coefficient was selected.

### 2.7 Static interaction fingerprint ranking

Static interaction fingerprints were generated from the prepared protein-ligand complexes using the Schrödinger structural interaction fingerprint (SIFt) implementation adapted to residue-level feature encoding [17]. Each docked pose and its reference complex were represented by a set of residue interaction features. A feature consisted of the hERG residue number paired with an interaction type: contact, hydrogen-bond donor or acceptor, hydrophobic, polar, backbone, or aromatic.

Because hERG is a homotetramer, equivalent interactions on different chains were treated as biologically equivalent. Interaction features were collapsed across chains, while the number of chains supporting each feature was retained as the chain-support count. Different interaction types at the same residue were kept as separate features.

Two quantities were used to compare a candidate pose with its reference fingerprint. The first, reference feature recovery (*R*), measures the fraction of reference residue interaction features that are present in the candidate pose:

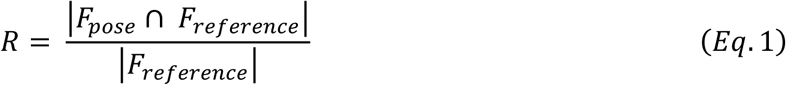

where *F* denotes the set of residue interaction features after chain collapse. The second quantity, cosine similarity (*C*), compares the chain-support vectors of the candidate and reference fingerprints:

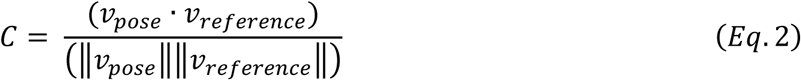

where each vector component corresponds to a residue interaction feature after chain collapse and stores the number of hERG chains supporting that feature. Thus, *R* measures recovery of the reference feature set, while *C* measures agreement in the multichain support pattern. The static interaction fingerprint score was calculated as an equally weighted average:

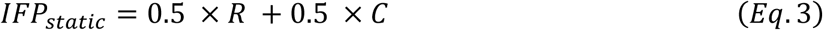

The highest static IFP criterion selected the pose with the largest *IFP_static_* score.

### 2.8 Composite static shape:IFP ranking

Shape and static IFP similarities were combined to test whether recovering both reference ligand volume and reference protein interactions improved pose selection. For each pose, the Tanimoto shape coefficient (*S*) and static IFP score (*IFP_static_*) were combined using three predefined weighting schemes: shape-favored (70:30), balanced (50:50), and IFP-favored (30:70):

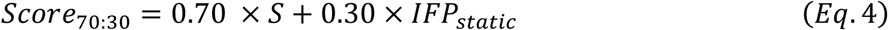

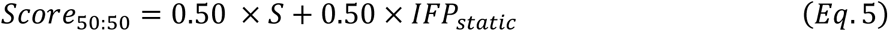

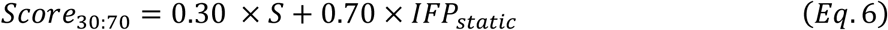

Six pose-selection criteria were evaluated for each docking system: lowest IFD score, highest static shape similarity, highest static IFP score, and the three composite shape:IFP scores. The pose ranked first by each criterion was advanced to MD evaluation.

### 2.9 Membrane system preparation and MD simulation

Selected complexes were prepared for MD using the Desmond System Builder [24]. Each hERG complex was embedded in a POPC lipid bilayer and solvated with TIP3P water in an orthorhombic simulation box. Systems were parameterized with the OPLS4 force field [18] and neutralized with counterions. The ligand protonation state and formal charge assigned during LigPrep were preserved.

Production MD simulations of 20 ns were performed in Desmond under the NPγT ensemble at 300 K and 1.01325 bar, using a Nosé-Hoover chain thermostat and a Martyna-Tobias-Klein barostat with default coupling parameters. The same protocol was used for docked poses and the corresponding reference complexes so that the pose quality descriptors were evaluated under matched simulation conditions.

### 2.10 Framewise MD interaction fingerprints

Protein-ligand interaction fingerprints were calculated frame-by-frame for each trajectory. Receptor and ligand atoms were taken from the coordinates of the same frame, allowing the interaction fingerprint to follow receptor motion during MD. Interactions were identified using the default SIFt definitions and distance criteria. For this analysis, a collapsed interaction feature was considered persistent when it was present in at least 50% of the analyzed frames.

### 2.11 MD-IFP similarity to reference trajectories

For each candidate pose, the MD interaction fingerprint was compared with the reference trajectory defined for that docking scenario. Interaction features were collapsed across chains as in the static IFP analysis, while chain-support counts were retained.

The MD-IFP similarity score (*IFPMD*) was defined as an equally weighted combination of three descriptors. The first, trajectory-level reference feature recovery (*RMD*), measured the fraction of reference interaction features recovered by the candidate trajectory:

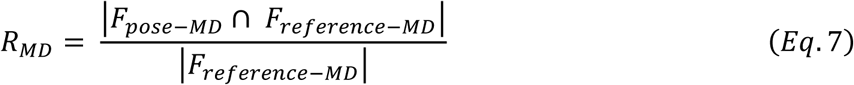

where *F_pose_*_−*MD*_ and *F_reference_*_−*MD*_ denote the sets of residue interaction features, after chain collapse, that were present in the candidate and reference trajectories, respectively.

The second descriptor, distinct chain count recovery (*CCMD*), measured how completely the candidate trajectory recovered the number of distinct hERG chains supporting each reference interaction feature. For each reference feature *f*, the chain count recovery was calculated as:

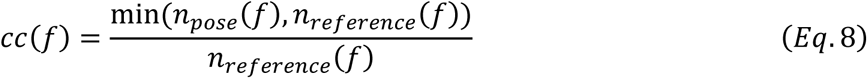

where *n_pose_*(*f*) and *n_reference_*(*f*) are the numbers of distinct hERG chains on which feature *f* was observed in the candidate and reference trajectories, respectively. The candidate chain-support count was capped at the reference value, so support beyond the reference did not receive additional credit. The overall MD chain count recovery descriptor was calculated as the mean of the per feature values across all interaction features present in the reference:

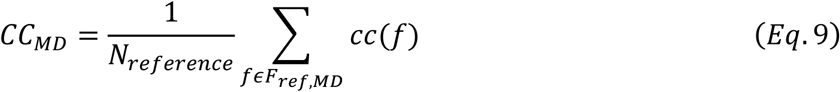

The third descriptor was the cosine similarity of the time-averaged chain-count vectors across collapsed interaction features (*CMD*), defined analogously to Eq. 2 using mean chain count vectors over the trajectory. The composite *IFPMD* score was then calculated as:

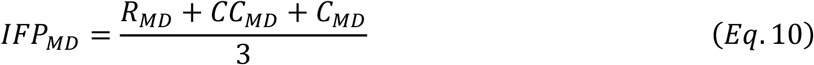

The three components capture complementary aspects of agreement during MD: *R_MD_* measures recovery of reference interaction features, *CC_MD_* measures recovery of chain support for those features, and *C_MD_* measures similarity in the mean chain count distribution over the trajectory.

### 2.12 Trajectory based structural descriptors

Ligand solvent accessible surface area (SASA), ligand RMSD, and protein-fitted ligand RMSD were calculated using the Schrödinger trajectory analysis tools. Mean ligand SASA over the production trajectory was used to assess ligand burial or exposure relative to the corresponding reference trajectory [25]. Ligand RMSD measured positional stability relative to the starting docked pose. Protein-fitted ligand RMSD was calculated after aligning the Cα atoms of the transmembrane segments to the starting complex and measured ligand displacement within the protein binding environment.

### 2.13 MD-quality score and system-level success

The four primary descriptors after MD were integrated into a composite MD-quality score. The weights were 40% for MD-IFP similarity, 40% for SASA agreement, 10% for ligand RMSD, and 10% for protein-fitted ligand RMSD. These weights were chosen to emphasize the two reference-relative descriptors, MD-IFP similarity and SASA agreement, while giving lower weights to the RMSD measures of positional stability. Each remaining component was implemented as a bounded linear penalty, so poor agreement reduced that component to zero without producing a negative contribution to the total score:

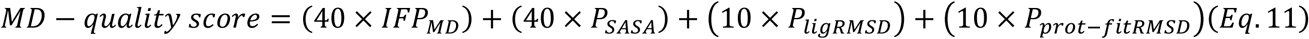

where:

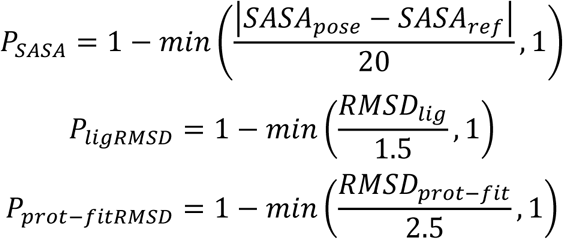

The cap values define failure for each descriptor. The ligand RMSD cap of 1.5 Å was selected as a stringent operational threshold for positional stability; a comparable low Å RMSD criterion has been reported in an IFD validation study [15]. The protein-fitted ligand RMSD cap of 2.5 Å was set wider to allow local receptor and ligand relaxation during MD while still penalizing large displacement within the binding environment. The SASA cap of 20 Å² was selected as a conservative operational threshold for changes in solvent exposure; classical solvent accessibility analyses indicate that residue-level ASA changes greater than 20 Å² are relatively uncommon [26]. The resulting MD-quality score is an operational score on a fixed 0-100 scale and was not statistically optimized.

To account for thermal drift in the reference trajectories, each pose score was also normalized to the reference score specific to that docking scenario:

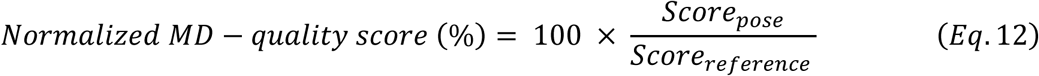

A ligand-template system was considered successful when at least one pose selected by the six predefined pose-selection criteria met both thresholds: an absolute MD-quality score of at least 75 and a normalized MD-quality score of at least 85% of the corresponding reference score. Using both thresholds reduced the chance that a poor pose would appear acceptable because the reference trajectory also drifted, and it also allowed modest thermal deviation in the reference trajectory. Success was assigned at the ligand-template system level. The composite descriptor is referred to as the MD-quality score, and its normalized form as the normalized MD-quality score.

## 3. Results

The dual criterion defined in Section 2.13, an absolute MD-quality score ≥ 75 together with a normalized MD-quality score ≥ 85% of the corresponding reference, was applied consistently across the retrospective ligand-template systems.

### 3.1 Chemical and structural basis of the proposed receptor environments

To define the chemical context represented by the available ligand-bound structures, the ligand set was characterized using physicochemical descriptors and the experimental reference complexes. Astemizole and pimozide occupied the low TPSA, high XLogP3-AA region, whereas E4031 occupied a more polar region and contained methanesulfonanilide functionality. Terfenadine grouped with astemizole and pimozide by these descriptors and substructure features, while dofetilide and ibutilide grouped with E4031 (Table 2). These group assignments describe the current structural set and provide the basis for the analyses below.

**Table 2.**
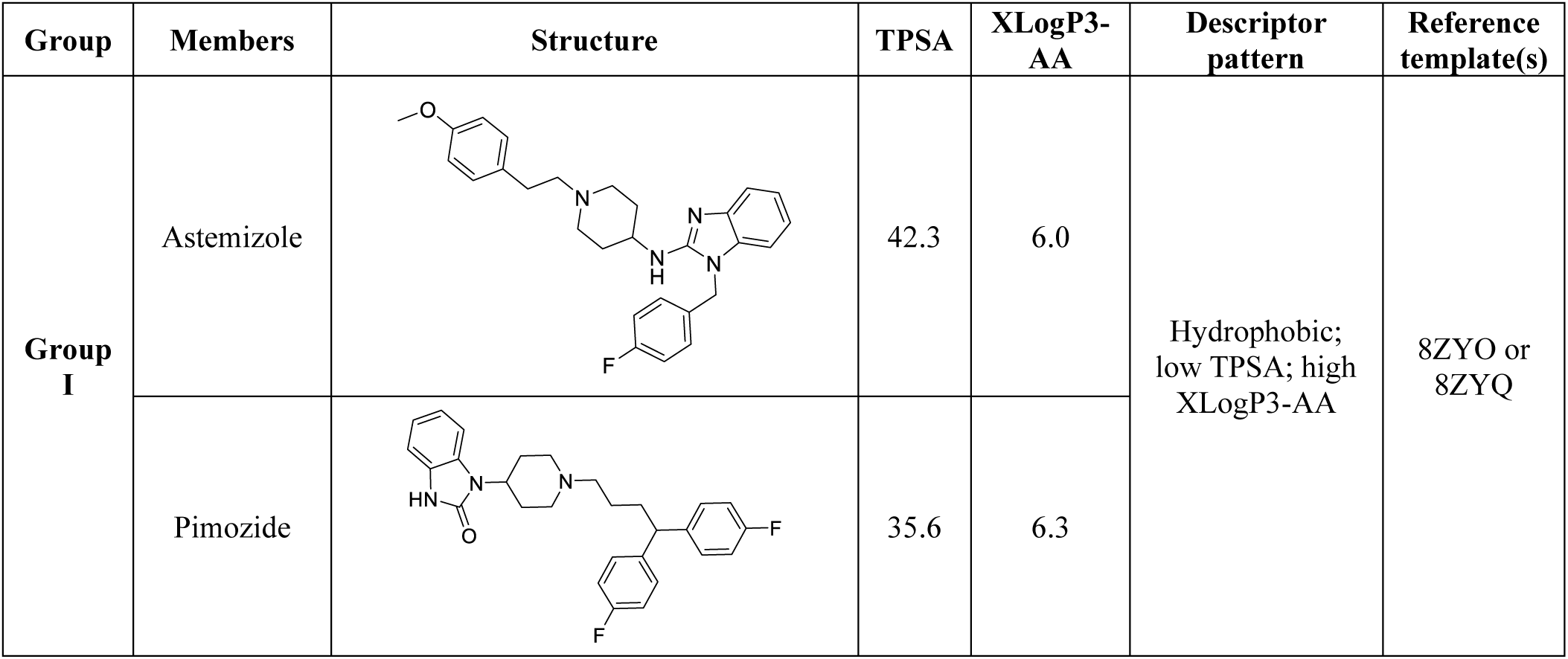
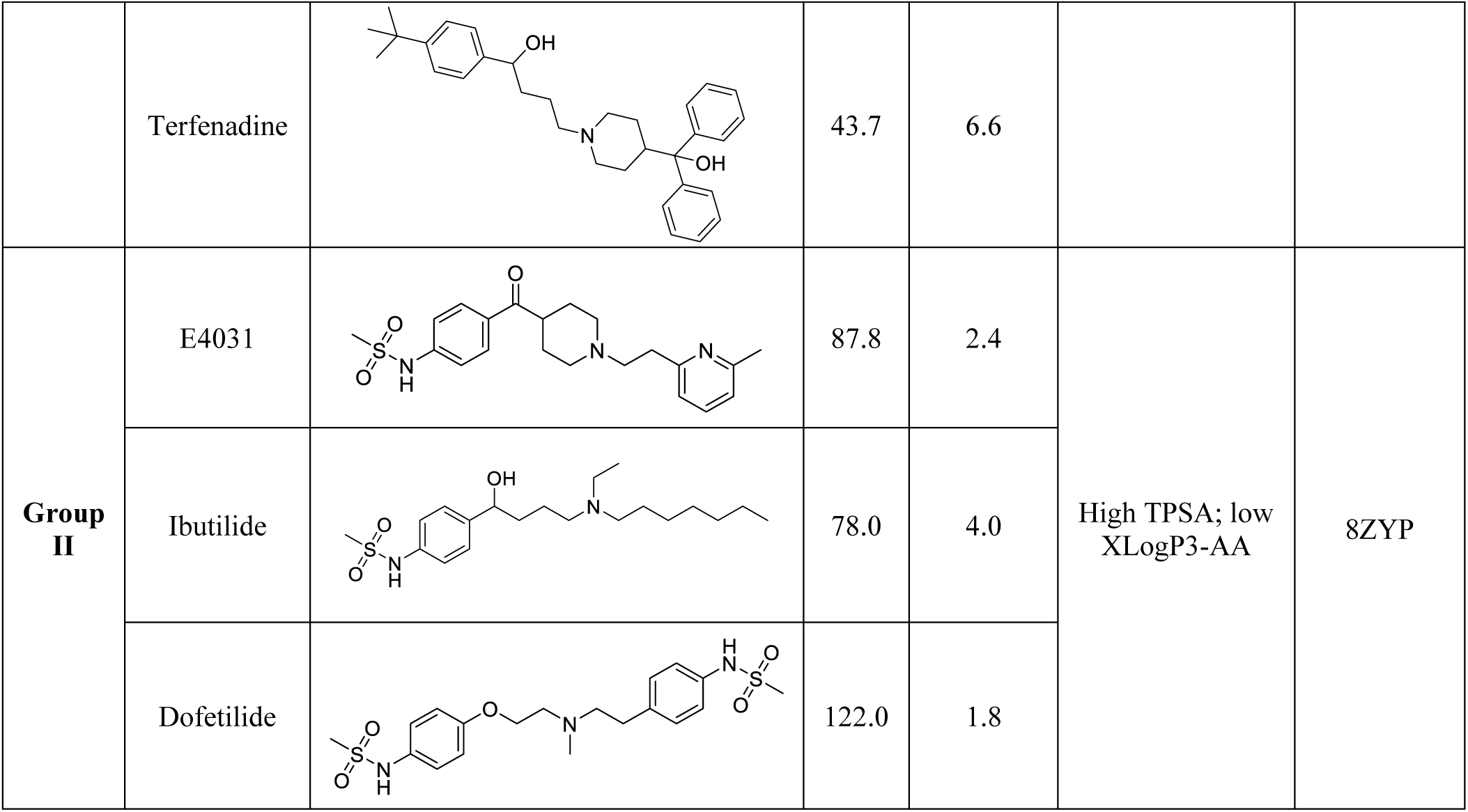
Study-defined group classification used to assign candidate receptor environments in this study. Physicochemical descriptors (TPSA and XLogP3-AA) were obtained from PubChem [**20**]

| Group | Members | Structure | TPSA | XLogP3-AA | Descriptor pattern | Reference template(s) |
| --- | --- | --- | --- | --- | --- | --- |
| Group I | Astemizole |  | 42.3 | 6.0 | Hydrophobic; low TPSA; high XLogP3-AA | 8ZYO or 8ZYQ |
|  | Pimozide |  | 35.6 | 6.3 |  |  |
|  | Terfenadine |  | 43.7 | 6.6 |  |  |
| **Group II** | E4031 |  | 87.8 | 2.4 | High TPSA; low XLogP3-AA | 8ZYP |
|  | Ibutilide |  | 78.0 | 4.0 |  |  |
|  | Dofetilide |  | 122.0 | 1.8 |  |  |

Group I comprised astemizole, pimozide, and terfenadine. Their TPSA values were below 60 Å² and their XLogP3-AA values exceeded 5. The astemizole- and pimozide-bound cryo-EM complexes (8ZYO and 8ZYQ) place large hydrophobic and aromatic ligand surfaces within the inner cavity and include contacts involving S624 and the S6 aromatic region, including Y652 and F656 [11]. Group II comprised E4031, dofetilide, and ibutilide. In the current ligand set, this region was characterized approximately by TPSA >70 Å² and XLogP3-AA <4.5, together with methanesulfonanilide, methanesulfonamide, or related polar functionality. E4031-bound 8ZYP provided the experimental structural reference for this region and included contacts involving S624, nearby residues in the pore helix and selectivity filter base, and S6 cavity residues [11]. The separation of the two groups in descriptor space is shown in Fig. 2.

**Fig. 2.**
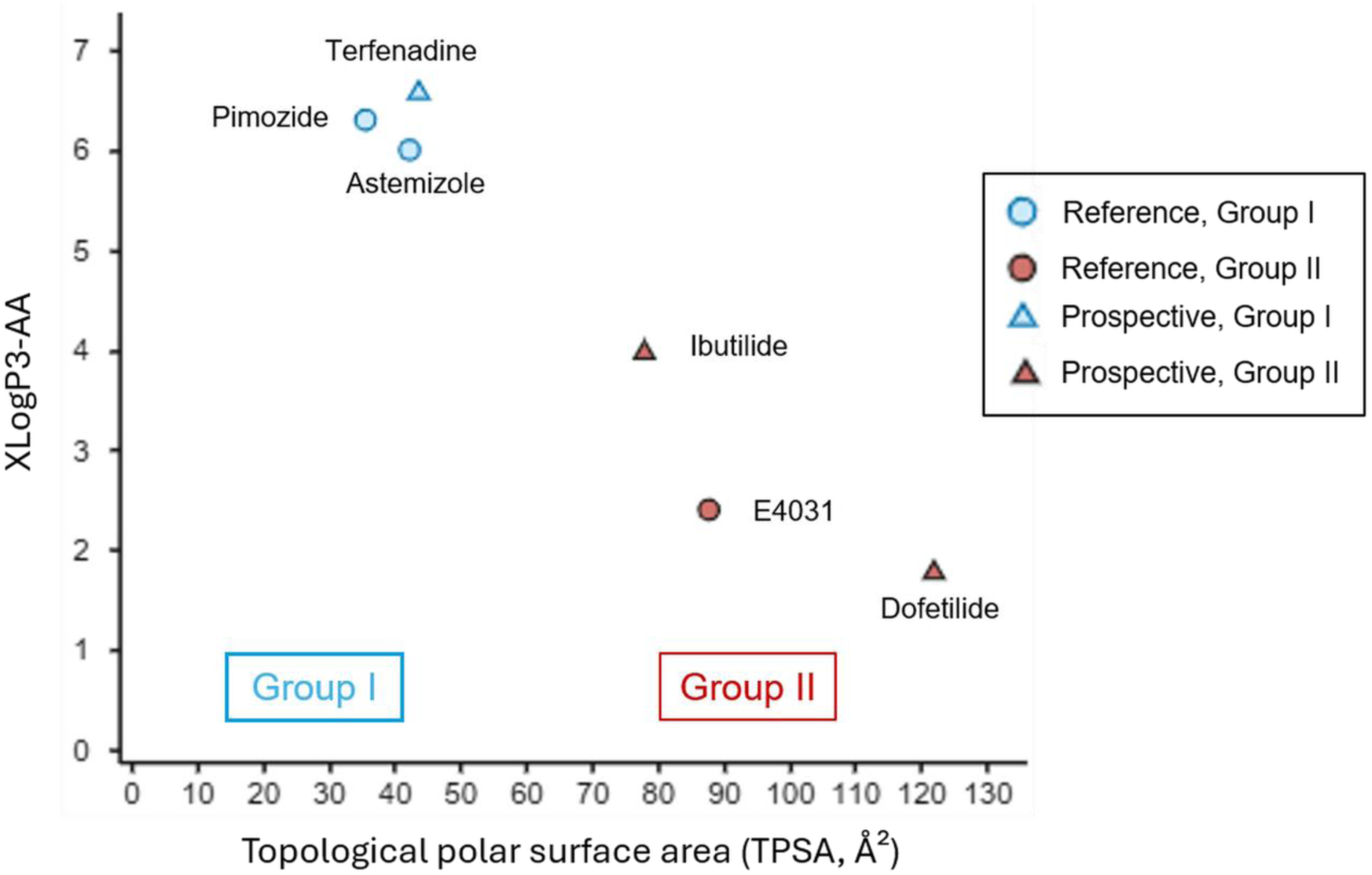
Distribution of Group I and Group II hERG blockers in TPSA-XLogP3-AA descriptor space. Circles represent cryo-EM reference ligands and triangles represent prospective ligands

For the analyses below, 8ZYO and 8ZYQ were designated Group I reference environments and 8ZYP was designated the Group II reference environment. A system was classified as group-matched when the docked ligand and the ligand defining the receptor template belonged to the same group defined in this study, and group-mismatched otherwise. This classification was used retrospectively to examine whether chemically similar ligand-template pairs were associated with more favorable cross-docking outcomes. Apo 8ZYN was analyzed separately as the receptor state control.

### 3.2 Can the proposed receptor environments support reliable pose generation?

#### 3.2.1 Self-docking: effect of receptor flexibility

Self-docking was used to compare pose generation when both the receptor environment and the experimental ligand orientation were known. Astemizole, pimozide, and E4031 were docked back into 8ZYO, 8ZYQ, and 8ZYP after removal of the resolved ligand, using rigid Glide and IFD. Representative poses from each protocol were advanced to MD and compared using the same descriptors after simulation (Table 3).

**Table 3.** Best self-docking performance of rigid Glide and induced-fit docking after MD. MD-q. = MD-quality score (max = 100); |ΔSASA| = deviation of mean ligand SASA from the cognate cryo-EM reference.

| Ligand-template system | Docking protocol | MD-IFP | SASA mean (Å<sup>2</sup>) | Ligand RMSD (Å) | Protein-fitted ligand RMSD (Å) | \|ΔSASA\| (Å<sup>2</sup>) | MD-q. |
| --- | --- | --- | --- | --- | --- | --- | --- |
| Astemizole-8ZYO | IFD | 0.958 | 13.59 | 0.73 | 1.39 | 1.93 | 84.0 |
| Astemizole-8ZYO | Glide | 0.960 | 16.69 | 0.88 | 1.77 | 5.03 | 75.4 |
| Pimozide-8ZYQ | IFD | 0.881 | 4.71 | 0.42 | 1.68 | 0.14 | 85.4 |
| Pimozide-8ZYQ | Glide | 0.870 | 13.62 | 0.94 | 1.93 | 9.05 | 62.7 |
| E4031-8ZYP | IFD | 0.866 | 17.03 | 0.42 | 0.94 | 9.62 | 68.8 |
| E4031-8ZYP | Glide | 0.530 | 70.93 | 0.77 | 1.96 | 44.28 | 28.2 |

IFD produced higher MD-quality scores than rigid Glide in all three self-docking systems. For astemizole-8ZYO, IFD and Glide showed comparable MD-IFP similarity (0.958 and 0.960), while the pose selected from IFD had a higher MD-quality score (84.0 versus 75.4), smaller SASA deviation from the cognate reference, and lower ligand RMSD. For pimozide-8ZYQ, the IFD pose had a higher MD-quality score (85.4 versus 62.7), lower ligand RMSD (0.42 versus 0.94 Å), and ligand SASA closer to the cognate reference. For E4031-8ZYP, IFD produced an MD-quality score of 68.8 compared with 28.2 for rigid Glide; the rigid docking pose showed greater solvent exposure (70.93 Å² versus 26.65 Å² for the cognate reference) and lower MD-IFP similarity (0.530 versus 0.866). IFD was therefore used for pose generation in the remaining docking experiments. The IFD poses and the corresponding cryo-EM reference poses are superposed in Fig. 3.

**Fig. 3.**
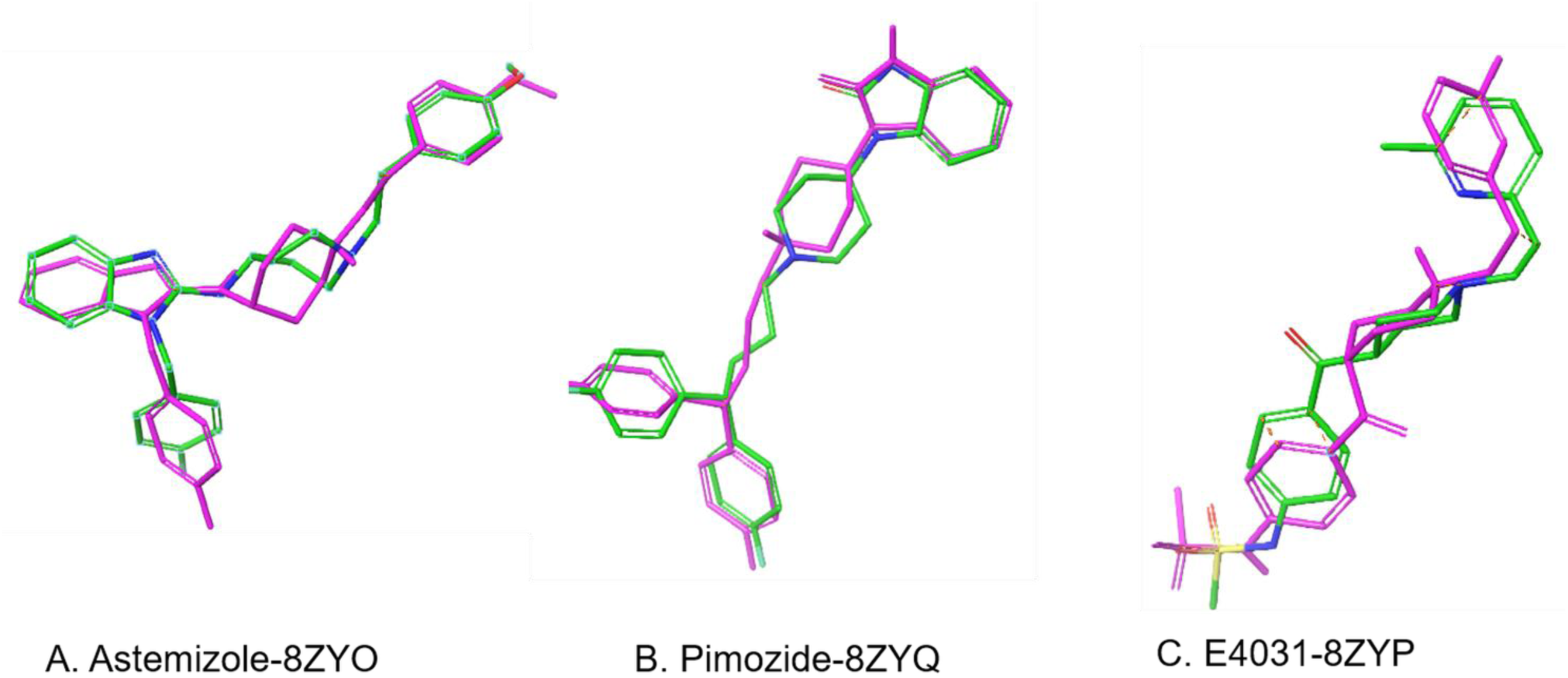
Self-docking superpositions of the three reference hERG inhibitors. Selected pre-MD IFD poses are shown in magenta and the corresponding cryo-EM reference poses in green: (A) astemizole-8ZYO, (B) pimozide-8ZYQ, and (C) E4031-8ZYP

#### 3.2.2 Apo hERG as a receptor state control

The inhibitor-free structure was evaluated as an alternative starting state. Astemizole, pimozide, and E4031 were each docked into apo 8ZYN, all six pose-selection criteria were applied, and the selected poses were advanced to MD. The maximum MD-quality scores were 44.6 for astemizole-8ZYN, 42.1 for pimozide-8ZYN, and 35.4 for E4031-8ZYN. All three values were below the predefined cutoff of 75 (Table 4).

**Table 4.** Apo state docking outcomes for the three reference inhibitors. All three systems fail the dual MD-quality criterion (raw ≥ 75 and normalized ≥ 85% of cognate reference MD-quality score)

| Ligand-template system | Max raw MD-quality | Normalized MD-quality (% of reference score) | Cognate reference MD-quality score | Outcome |
| --- | --- | --- | --- | --- |
| Astemizole-8ZYN apo | 44.6 | 51.7% | 86.3 | FAIL |
| Pimozide-8ZYN apo | 42.1 | 47.1% | 89.4 | FAIL |
| E4031-8ZYN apo | 35.4 | 40.8% | 86.7 | FAIL |

For all three apo systems, mean ligand SASA exceeded the corresponding ligand-bound reference by approximately 34 Å² for astemizole, 63 Å² for pimozide, and 50 Å² for E4031, exceeding the 20 Å² cap in every case. Ligand RMSD and MD-IFP similarity varied among the systems, but none met the dual criterion. Apo 8ZYN was therefore retained as a receptor state control and was not used as a primary template in the transferability analysis.

### 3.3 Does chemical matching between ligand and template predict transferability across hERG structures?

#### 3.3.1 Ligand-referenced cross-docking

The ligand-bound structures were next evaluated by cross-docking. In ligand-referenced cross-docking, each reference inhibitor was docked into the other two ligand-bound templates and evaluated against its own experimentally resolved hERG pose. Six systems were examined: astemizole in 8ZYP and 8ZYQ; pimozide in 8ZYO and 8ZYP; and E4031 in 8ZYO and 8ZYQ.

The best MD-quality score varied across ligand and template combinations. Pimozide-8ZYO and astemizole-8ZYQ reached scores of 83.1 and 81.5, respectively. E4031-8ZYQ and E4031-8ZYO reached 61.6 and 69.7, astemizole-8ZYP reached 74.0, and pimozide-8ZYP reached 41.2. For each inhibitor, changing the ligand-bound receptor template changed the outcome after MD.

#### 3.3.2 Template-referenced cross-docking without a cognate ligand pose

The cross-docking analysis was then repeated under the template-referenced condition. The ligand already resolved in each receptor template served as the structural reference. Candidate poses were ranked using the template-bound ligand and its interaction fingerprint. The cognate pose of the docked ligand was not used in this analysis.

Template-referenced cross-docking produced the same broad separation observed in the ligand-referenced analysis (Table 5). Pimozide-8ZYO and astemizole-8ZYQ passed the dual MD-quality criterion, with raw scores of 81.7 and 84.5 and normalized scores of 91.5% and 98.0%, respectively. Astemizole-8ZYP had a raw score of 74.0 and a normalized score of 85.7%, so it failed because the raw score was below 75. E4031-8ZYO, E4031-8ZYQ, and pimozide-8ZYP produced MD-quality scores of 49.6, 67.9, and 51.1, respectively. The two within Group I cross-docking pairs passed, while the group-mismatched pairs remained below the dual criterion.

**Table 5.** Template-referenced cross-docking outcomes. Values report the best MD-quality score for each cross-docked ligand-template pair.

| Docked ligand | Template | Reference complex | MD-quality score | % of reference score | Outcome (dual criterion) |
| --- | --- | --- | --- | --- | --- |
| Pimozide | 8ZYO | Astemizole-8ZYO | 81.7 | 91.5% | Pass |
| Astemizole | 8ZYQ | Pimozide-8ZYQ | 84.5 | 98.0% | Pass |
| Astemizole | 8ZYP | E4031-8ZYP | 74.0 | 85.7% | Fail (raw < 75) |
| E4031 | 8ZYO | Astemizole-8ZYO | 49.6 | 57.2% | Fail |
| E4031 | 8ZYQ | Pimozide-8ZYQ | 67.9 | 78.3% | Fail |
| Pimozide | 8ZYP | E4031-8ZYP | 51.1 | 57.2% | Fail |

#### 3.3.3 Group match status and system-level behavior

The cross-docking outcomes were then tabulated according to the group assignments described in Section 3.1. Across the retrospective ligand-bound systems, six of seven group-matched systems passed the dual criterion, while all eight group-mismatched systems failed (Table 6). The only group-matched failure was E4031 self-docking in 8ZYP, which remained below the system-level criterion because of reduced SASA agreement despite near-reference positional behavior.

**Table 6.** System level outcomes according to ligand-template group-match status.

| System category | Number of systems | Pass (dual criterion) | Fail | Success rate |
| --- | --- | --- | --- | --- |
| Group-matched ligand-bound | 7 | 6 | 1 | 86% |
| Group-mismatched ligand-bound | 8 | 0 | 8 | 0% |
| Apo 8ZYN | 3 | 0 | 3 | 0% |

### 3.4 Which reference information best prioritizes poses within compatible templates?

Each IFD calculation generated multiple candidate poses, so the six predefined static selection criteria were compared within the retrospective group-matched systems for which experimental reference information was available. The Group I retrospective set contained six self-docking or cross-docking systems. E4031-8ZYP was the only retrospective Group II ligand-bound system because 8ZYP was the only Group II experimental reference in the current structural set. Criterion results and static descriptors are provided in Supplementary Tables S1 and S2.

For Group I, highest static shape similarity selected a pose that passed the dual criterion in all six retrospective systems (6/6), with raw MD-quality scores of 77.4-85.4 and normalized scores of 86.6%-98.0% of the cognate reference. Each of the other five criteria failed in at least one Group I system. The shape-selected poses retained Y652 interactions across multiple subunits together with F656 contacts. Highest static shape similarity was therefore carried forward as the Group I pose ranking criterion for the prospective analysis.

For Group II, no pose-selection criterion brought the single retrospective E4031-8ZYP system above the dual MD-quality threshold. The highest raw score was 68.8 and was shared by the highest static shape, 70:30, 50:50, and 30:70 criteria, which converged on the same pose. The pose selected by the 30:70 criterion showed persistent S624, V625, and Y652 interactions with transient T623 sampling during MD. Because E4031-8ZYP was the only retrospective Group II ligand-bound system, the 30:70 IFP-favored composite was carried forward as a provisional Group II criterion.

The lowest IFD score did not reach the normalized score threshold in any group-matched retrospective system. The prospective analyses therefore used the group-specific reference-informed criteria identified above as the primary selection criteria. The lowest IFD score was retained as a comparator. The normalized MD-quality scores for each criterion are summarized in Fig. 4.

**Fig. 4.**
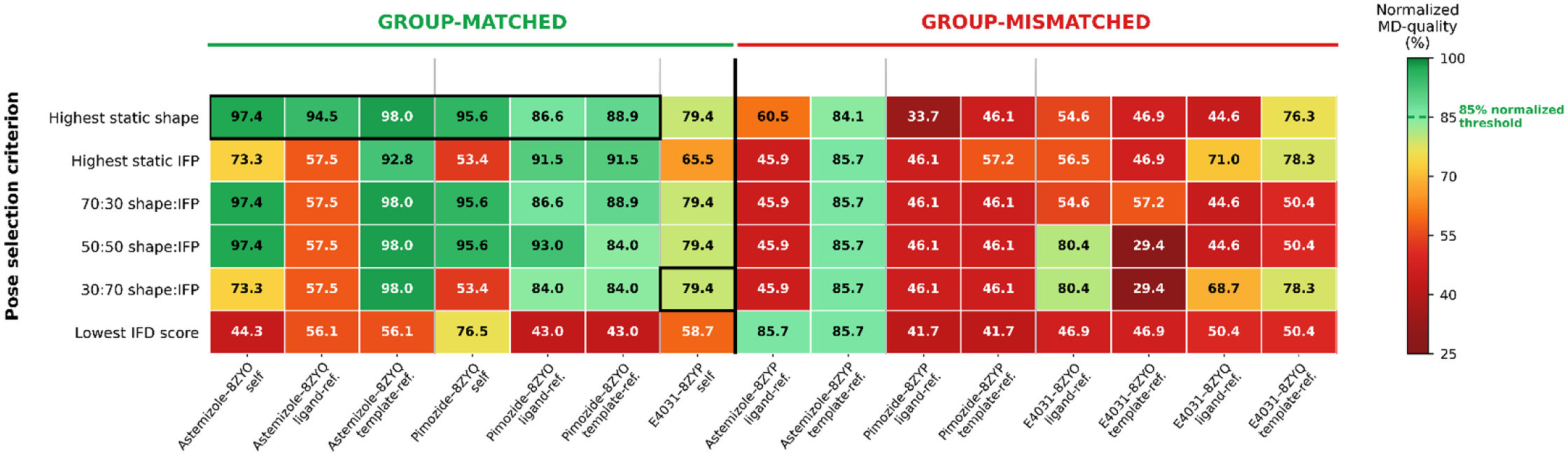
Performance of pose-selection criteria across ligand-bound hERG docking systems. Cell values are normalized MD-quality score (% of cognate reference MD-quality score). Boxed cells indicate the group-specific strategy carried forward for prospective evaluation

### 3.5 Prospective test of the complete framework

Dofetilide, ibutilide, and terfenadine were used to evaluate the complete framework for ligands that were not among the three experimental cryo-EM reference ligands. Each compound was classified using physicochemical descriptors and substructure information available before docking, then docked into all three ligand-bound receptor templates. The ranking criterion associated with its group was used for the primary assignment. Alternative template comparisons were retained for all three compounds. Static shape similarity and static IFP scores for the selected poses across all three ligand-bound receptor templates are provided in Supplementary Table S3. MD descriptors and residue contacts were recorded for the selected poses; comparison with independent residue-level literature is considered in the Discussion.

#### 3.5.1 Dofetilide (Group II)

Dofetilide contains two methanesulfonanilide groups and an ether linker. Its TPSA (122 Å²), XLogP3-AA (1.8), and methanesulfonanilide pharmacophore placed it in Group II. It was evaluated primarily in E4031-bound 8ZYP using the provisional 30:70 IFP-favored composite, with 8ZYO and 8ZYQ retained for alternative template comparison.

In 8ZYP, the 30:70 composite and highest static IFP criterion converged on the same pose, while the lowest IFD score and highest static shape criteria selected different poses. The pose selected by the 30:70 criterion recovered the full E4031 reference feature set after chain collapse and occupied the cavity region spanning S624/Y652/G648/L622 (Fig. 5). During MD, the pose sampled S624, V625, G648, T623, and Y652, with the strongest dynamic contacts at Y652, V625, and S624. Mean ligand SASA was 19.16 ± 5.93 Å², ligand RMSD was 0.77 ± 0.27 Å, and protein-fitted ligand RMSD was 1.27 ± 0.29 Å. Among the three receptor structures tested, the 8ZYP pose showed the strongest combined agreement with the template reference and the MD descriptors for dofetilide.

**Fig. 5.**
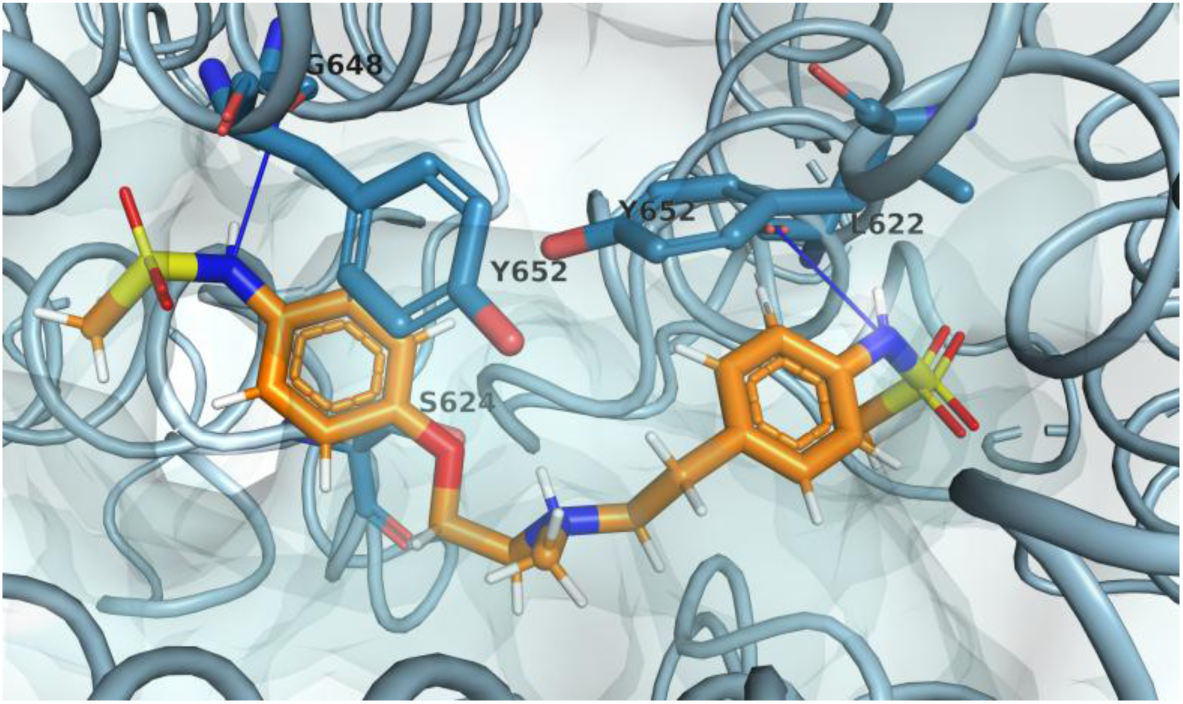
Representative MD frame for the dofetilide-8ZYP pose selected using the 30:70 IFP-favored composite. Dofetilide is shown in orange within the E4031-bound hERG pore cavity

#### 3.5.2 Ibutilide (Group II)

Ibutilide contains a single methanesulfonanilide group, a secondary alcohol, and an aliphatic chain. Its TPSA (78 Å²), XLogP3-AA (4.0), and methanesulfonanilide pharmacophore placed it in Group II. Ibutilide was evaluated primarily in E4031-bound 8ZYP using the provisional 30:70 IFP-favored composite, with 8ZYO and 8ZYQ retained for alternative template comparison.

In 8ZYP, the 30:70 composite and highest static IFP criterion converged on the same selected pose. During MD, the pose sampled Y652 aromatic and hydrophobic contacts across multiple subunits, S624 hydrogen bonding from the methanesulfonanilide region, V625 hydrophobic contact across three subunits, and T623 contact (Fig. 6). Mean ligand SASA was 19.92 ± 9.60 Å², ligand RMSD was 1.14 ± 0.27 Å, and protein-fitted ligand RMSD was 2.00 ± 0.35 Å. Ibutilide also produced cavity poses in 8ZYO and 8ZYQ with S624, Y652, and F656 interactions. Poses from the alternative templates did not show the same combined S624/V625/Y652 interaction pattern as the 8ZYP-selected pose.

**Fig. 6.**
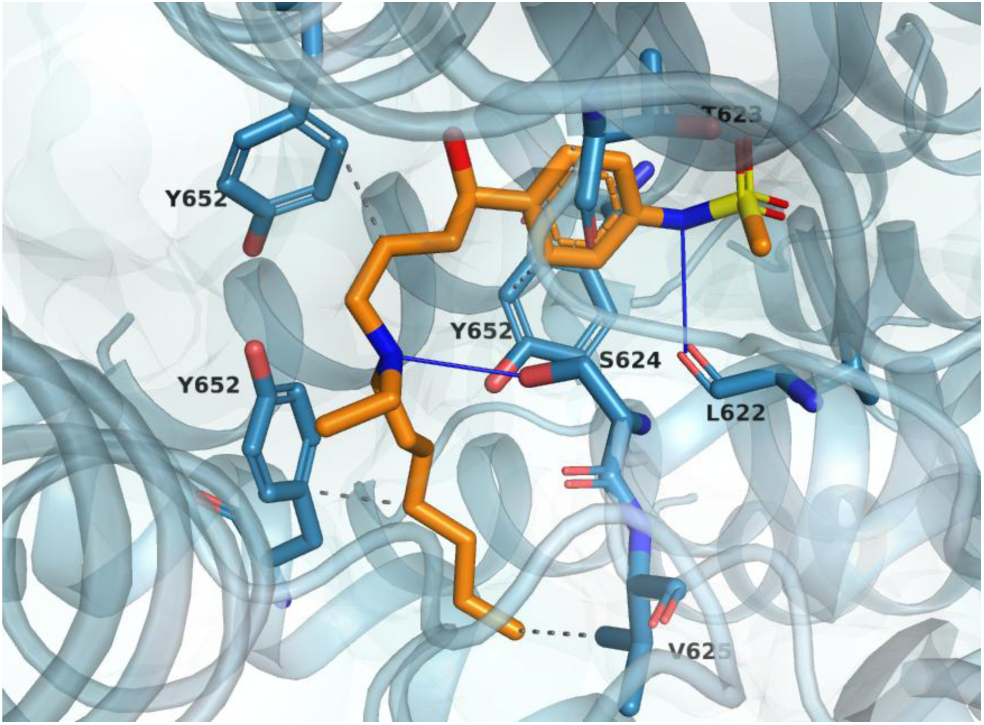
Representative MD frame for the ibutilide-8ZYP pose selected using the 30:70 IFP-favored composite. Ibutilide is shown in orange within the E4031-bound hERG pore cavity

#### 3.5.3 Terfenadine (Group I)

Terfenadine has a diphenyl scaffold bearing a tert-butyl substituent and a hydroxymethylpiperidinol head, and it lacks a methanesulfonanilide pharmacophore. Its TPSA (43.7 Å²), XLogP3-AA (6.6), and hydrophobic scaffold placed it in Group I. Terfenadine was evaluated primarily in the Group I receptor environments using highest static shape similarity, while all three ligand-bound templates were retained for comparison.

In 8ZYO, highest static shape similarity selected a pose that retained Y652 and F656 aromatic and hydrophobic contacts across multiple subunits together with T623 and S624 hydrogen bonding during the 20 ns MD trajectory (Fig. 7). V625 and G648 were not sustained contacts. The pose remained tightly buried, with mean ligand SASA of 11.81 Å², ligand RMSD of 0.74 Å, and protein-fitted ligand RMSD of 1.07 Å; its mean SASA closely matched the astemizole reference value of 11.66 Å². Terfenadine in 8ZYQ produced a less buried pose, with mean ligand SASA of 36.67 Å² compared with 4.57 Å² for the pimozide reference. Docking into 8ZYP produced poses with mean ligand SASA greater than 80 Å². Among the three receptor structures tested, 8ZYO provided the strongest support after MD for the selected terfenadine pose.

**Fig. 7.**
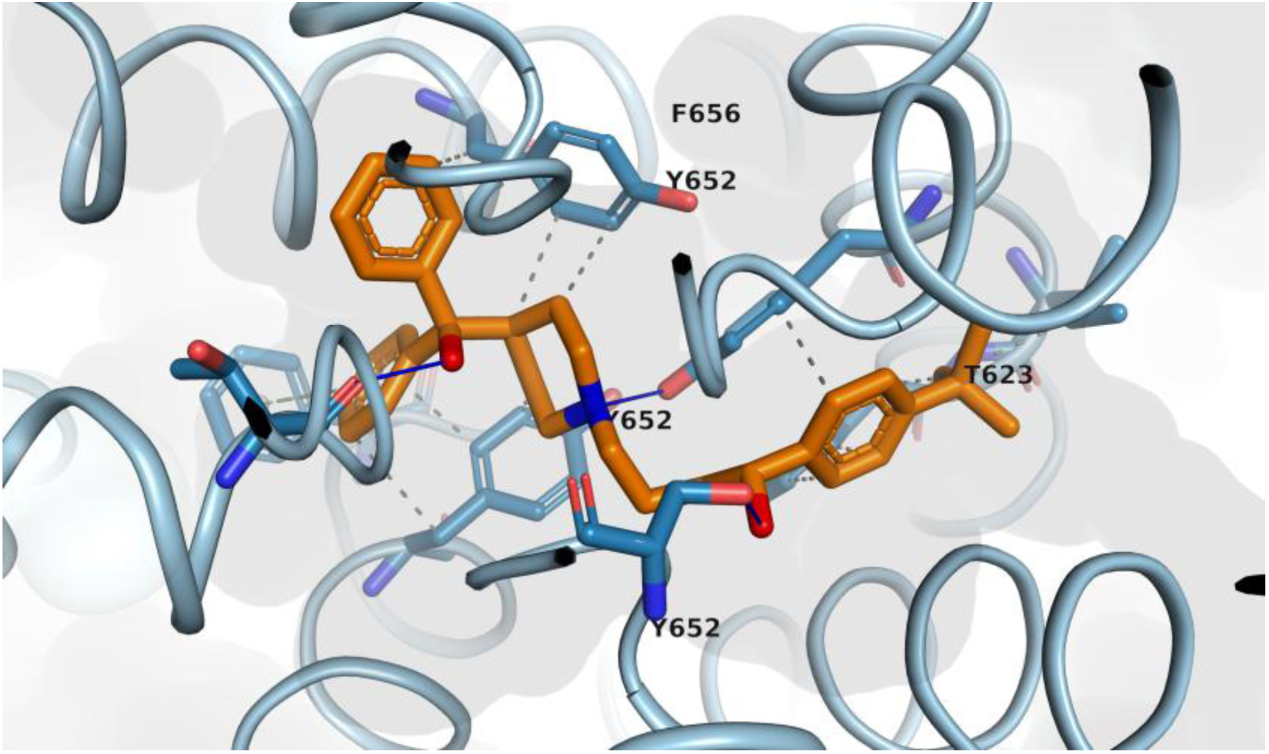
Representative MD frame for the terfenadine-8ZYO pose selected using highest static shape similarity. Terfenadine is shown in orange within the astemizole-bound hERG pore cavity

The prospective outcomes are summarized in Table 7. Dofetilide and ibutilide were assigned to E4031-bound 8ZYP using the provisional 30:70 IFP-favored composite, while terfenadine was assigned to astemizole-bound 8ZYO using highest static shape similarity. The selected poses remained within the hERG cavity during the analyzed trajectories and retained the residue-contact patterns reported above. Alternative template calculations gave lower overall support for the primary assignments, although some ligands also produced cavity poses in other receptor environments.

**Table 7.** Summary of prospective ligand-binding results.

| Ligand | Group | Template | Pose selection strategy | Static shape | Static IFP | Mean SASA (Å<sup>2</sup>) | MD residue contacts for selected pose |
| --- | --- | --- | --- | --- | --- | --- | --- |
| Dofetilide | II | 8ZYP | 30:70 IFP-favored composite | 0.554 | 0.961 | 19.16 | S624, V625, G648, T623, Y652 |
| Ibutilide | II | 8ZYP | 30:70 IFP-favored composite | 0.435 | 0.948 | 19.92 | S624, V625, Y652, T623 |
| Terfenadine | I | 8ZYO | Highest static shape | 0.660 | 0.784 | 11.81 | T623, S624, Y652, F656; V625 and G648 not sustained |

## 4. Discussion

### 4.1 Ligand-bound hERG structures as ligand-conditioned receptor microenvironments

The template comparison suggests that each ligand-bound hERG structure captures a distinct local receptor environment. Miyashita and co-workers resolved astemizole, pimozide, and E4031 in the same general inner cavity, but the local organization of residues in the pore helix, selectivity filter base, and S6 region differed with the bound ligand [11]. Sun, Wang, and Shen later reported high overall structural similarity among the apo and ligand-bound hERG structures, with pairwise Cα RMSDs below 1 Å, together with substantial flexibility of the Y652 side chains in the binding pocket [12]. Our docking results show that these local differences can affect how well another ligand is accommodated even when the overall structures are similar. The available complexes are therefore more usefully viewed as experimentally observed, ligand-conditioned receptor microenvironments than as equivalent copies of a single open-state template. The poor performance of apo 8ZYN under the present protocol further indicates that the local organization of the starting structure matters for prospective pose selection.

### 4.2 Physicochemical basis of ligand-template compatibility

The chemical groups defined in this study offer one possible explanation for the observed template dependence and describe the current applicability domain. Group I occupies a region of low polarity and high lipophilicity (approximately TPSA <60 Å² and XLogP3-AA >5) with extensive hydrophobic and aromatic surface area. Group II occupies a more polar region (approximately TPSA >70 Å² and XLogP3-AA <4.5) and contains methanesulfonanilide, methanesulfonamide, or related polar functionality. These differences are consistent with the distinct cavity environments represented by the astemizole/pimozide structures and E4031. The group-matched and group-mismatched results suggest that ligand chemistry can provide useful prior information when choosing among the available ligand-bound hERG templates. The boundaries should nevertheless be treated as approximate descriptors of the present dataset rather than fixed decision thresholds for all hERG chemotypes.

### 4.3 Pose generation and pose ranking are separate problems

The self-docking results distinguish pose generation from pose ranking. IFD improved the generated poses compared with rigid Glide, consistent with local receptor relaxation during docking [15]. However, the IFD score did not consistently identify the pose with the strongest support after MD. Broader docking benchmarks have reached a similar conclusion: generating a pose close to the experimental geometry and ranking that pose first are separate problems [13]. The ligand-bound cryo-EM complexes therefore contribute in two ways in this study. They provide receptor coordinates for pose generation and experimental ligand shape and interaction patterns for pose prioritization. This separation avoids asking a single empirical docking score to solve both sampling and final pose discrimination.

### 4.4 Why shape and interaction criteria may differ by ligand type

The static ranking criteria behaved differently across the two chemical regions. In Group I, the broad hydrophobic and aromatic character of the ligands may make overall three-dimensional ligand shape particularly informative, because related ligand conformations can retain similar global shape despite differences in their specific protein interactions or symmetry-equivalent binding orientations. The more polar E4031-associated Group II hypothesis places greater emphasis on directional and residue-specific interactions near S624, V625, T623, and adjacent pore-cavity residues, which may explain the value of an interaction-weighted criterion. This explanation remains preliminary because Group II is represented retrospectively by only one experimental ligand-bound complex, whereas Group I is supported by two reference structures and several self-docking and cross-docking combinations. The current data support evaluating different sources of reference information by ligand type while additional Group II structures are still needed.

### 4.5 Role of MD in pose assessment

MD provides a third layer of evidence rather than another docking score. It was used to examine how the selected poses behaved after relaxation in a membrane environment. Static shape and IFP comparisons evaluate the starting docked geometry against a structural reference, while MD evaluates whether the pose remains coherent as the protein, ligand, membrane, solvent, and ions relax. The MD-quality score combines interaction recovery, ligand burial, and positional stability into an operational pose assessment metric. The score does not estimate binding free energy or affinity and cannot establish a unique experimental binding mode. A 20 ns trajectory also cannot sample binding and unbinding kinetics, drug trapping, gating transitions, or long-timescale conformational changes. Within these limits, loss of expected contacts, marked solvent exposure, or displacement from the cavity provides evidence against a pose even when its initial static score is favorable.

### 4.6 Prospective interpretation and relation to experimental residue evidence

The prospective compounds show how the workflow behaves when a ligand-specific hERG structure is unavailable. For dofetilide, the 8ZYP-selected pose sampled S624, V625, T623, G648, and Y652, overlapping with residue regions implicated experimentally for dofetilide/E4031-like block [5, 27] . The ibutilide pose selected in 8ZYP sampled S624, V625, T623, and Y652, consistent with residues implicated by mutagenesis and structure-activity studies of clofilium and ibutilide analogs [6, 28] . For terfenadine, the 8ZYO-selected pose retained T623, S624, Y652, and F656, while V625 and G648 were not sustained; this pattern qualitatively matches the residue dependence reported for terfenadine/cisapride block [29]. These comparisons provide external mechanistic consistency rather than structural validation. Mutagenesis can reflect direct contact, allosteric coupling, altered channel state, or changes in accessibility. The prospective results therefore narrow the set of plausible binding hypotheses for these ligands.

### 4.7 Applicability domain, limitations, and future validation

The main limitation of the framework is the small structural reference set. Only three higher-resolution ligand-bound hERG complexes were available for this analysis, and the Group II branch is anchored retrospectively by E4031 alone. Intermediate or chemically distinct blockers may fall between the current chemical regions, so comparison across several ligand-bound templates is appropriate for such compounds. The receptor structures used here represent open hERG inner cavity conformations. The present analysis does not address binding to closed states, inactivation-dependent affinity, channel gating transitions, drug trapping, or binding and unbinding kinetics. The MD-quality weights and thresholds are operational choices for pose assessment, and the short simulations were designed to assess local structural consistency. They do not predict potency, binding affinity, state-dependent kinetics, or clinical proarrhythmic risk. Future evaluation should include additional ligand-bound hERG structures and chemically broader blockers, particularly compounds near the boundaries and at the extremes of the current descriptor space that also have independent residue-level experimental data. With broader structural coverage, the framework could develop into an expandable map linking ligand chemistry to experimentally observed receptor environments and pose-selection evidence.

## 5. Conclusion

This study evaluated a structure-based strategy for prospective hERG pose selection using experimentally resolved ligand-bound structures as receptor environments and structural references. The results show that receptor choice, pose generation, pose ranking, and dynamic assessment each contribute to uncertainty in prospective modeling. Within the current dataset, successful ligand transfer was concentrated in group-matched ligand-bound systems, while apo 8ZYN did not reproduce the behavior of the ligand-bound reference systems.

IFD improved pose generation relative to rigid Glide, but docking score alone was not sufficient for final pose prioritization. Reference informed shape similarity was most consistent across the retrospective Group I systems. For the E4031-associated Group II environment, the 30:70 IFP-favored composite was retained as a provisional strategy. Prospective application to dofetilide, ibutilide, and terfenadine produced mechanistically interpretable poses that remained coherent during short MD simulations and showed residue-contact patterns consistent with independent hERG literature.

The framework provides a structured approach for prioritizing binding-mode hypotheses. It is not intended to predict hERG potency or clinical risk. In practice, the receptor environment can be selected using ligand chemistry, candidate poses can be ranked with information beyond docking score, and the selected poses can be examined dynamically before mechanistic interpretation. Additional ligand-bound hERG structures will be needed to test and expand the current applicability domain. The same general strategy could also be tested for other proteins with flexible or ligand-dependent binding sites when suitable ligand-bound reference structures are available.

## Acknowledgements

The authors acknowledge the University of Connecticut Information Technology Services High Performance Computing (HPC) facility for providing the computational resources used in this study.

## Funding

This work was supported by QuantumCT.

## Competing Interests

The authors have no relevant financial or non-financial interests to disclose.

## Author Contributions

**PN:** Conceptualization, methodology, validation, formal analysis, investigation, data curation, visualization, writing – original draft, and writing – review & editing. **UO:** Conceptualization, supervision, and writing – review & editing. **SN:** Writing – review & editing, formal analysis, and visualization. **BC:** Supervision, project administration, funding acquisition, and writing – review & editing.

## Use of Artificial Intelligence

Generative AI tools, including Claude (Anthropic) and ChatGPT (OpenAI) were used for manuscript editing and for coding. It was not used to generate research data or results, and it is not an author.

## Data Availability

The data are available from the corresponding author upon reasonable request.

## Notes

### Competing Interest Statement

The authors have declared no competing interest.

