## Supporting Information for "A Structure-Based hERG Pose Selection Workflow Integrating Induced-Fit Docking, Shape Similarity, Interaction Fingerprints, and Molecular Dynamics"

Supplementary Fig. S1 summarizes the computational implementation of the hERG pose-selection analysis. Tables S1 and S2 provide the pose-selection criterion results and static descriptors for the group-matched retrospective ligand-bound systems reported in Section 3.4 of the manuscript. Table S3 provides static similarity scores for the prospective ligands across all three ligand-bound receptor templates evaluated in Section 3.5.

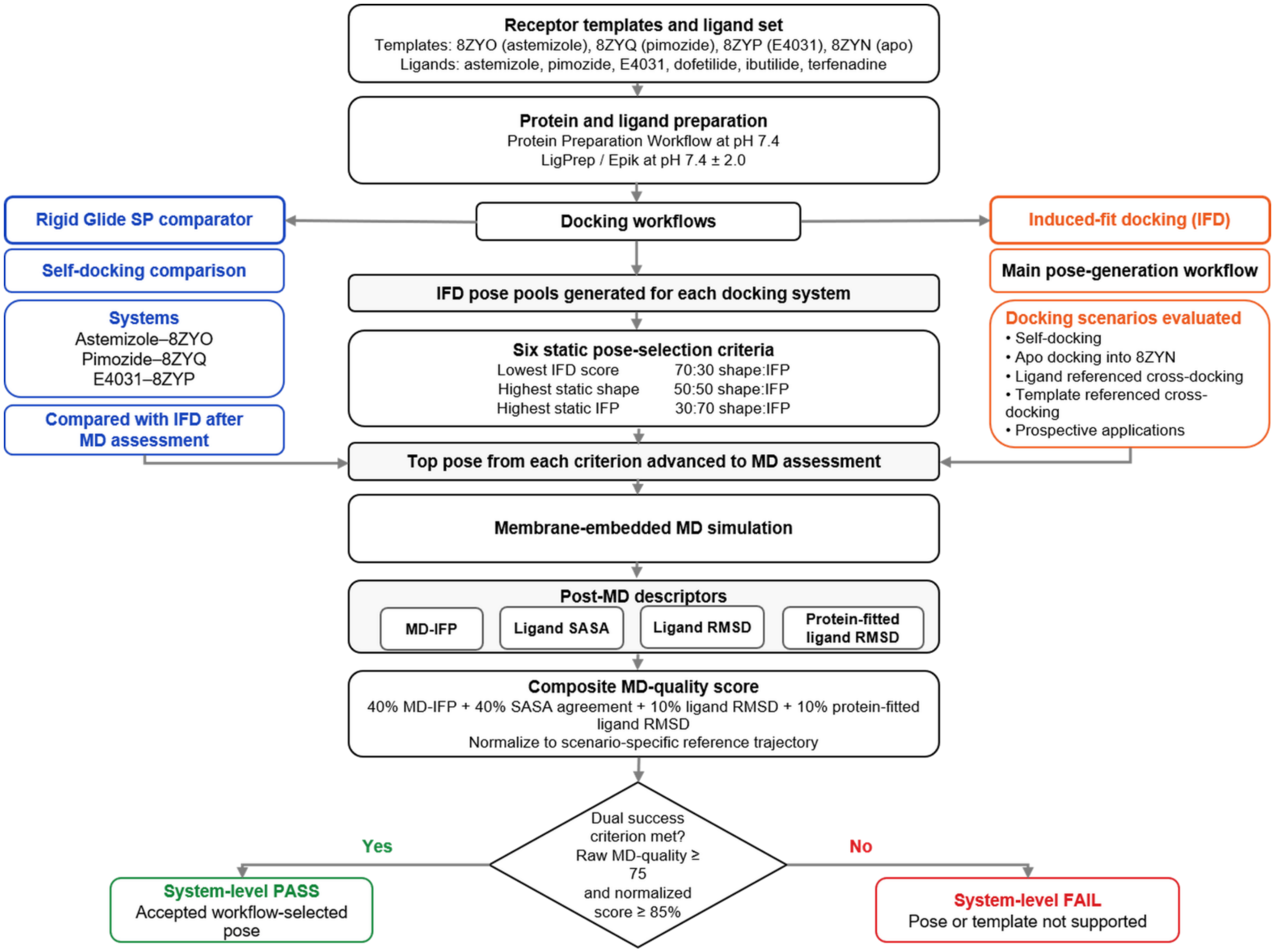


**Supplementary Fig. S1** Computational implementation of the hERG pose-selection analysis, integrating receptor-template selection, docking, static shape and interaction fingerprint rescoring, MD in a membrane environment, and pose quality assessment after MD

### Supplementary Table. S1 Pose-selection criterion performance across group-matched retrospective ligand-bound systems

| Pose-selection criterion | Astemizole–8ZYO Self-docking | Pimozide–8ZYQ Self-docking | Pimozide–8ZYO Ligand-ref. | Astemizole–8ZYQ Ligand-ref. | Pimozide–8ZYO Template-ref. | Astemizole–8ZYQ Template-ref. | E4031–8ZYP Self-docking |
|---|---|---|---|---|---|---|---|
| Highest static shape similarity | 97.4 | 95.6 | 86.6 | 94.5 | 88.9 | 98.0 | 79.4 |
| Highest static IFP score | 73.3 | 53.4 | 91.5 | 57.5 | 91.5 | 92.8 | 65.5 |
| 70:30 shape-favored composite | 97.4 | 95.6 | 86.6 | 57.5 | 88.9 | 98.0 | 79.4 |
| 50:50 balanced composite | 97.4 | 95.6 | 93.0 | 57.5 | 84.0 | 98.0 | 79.4 |
| 30:70 IFP-favored composite | 97.4 | 95.6 | 84.0 | 57.5 | 84.0 | 98.0 | 79.4 |
| Lowest IFD score | 44.3 | 76.5 | 43.0 | 56.1 | 43.0 | 56.1 | 58.7 |

Values are normalized MD-quality scores (% of the reference MD-quality score specific to each docking scenario). The dual criterion requires both an absolute MD-quality score ≥75 and a normalized MD-quality score ≥85%. The first six systems are Group I; E4031–8ZYP is Group II.

Ligand-ref., ligand-referenced cross-docking using the docked ligand's own cryo-EM reference; template-ref., template-referenced cross-docking using the template-bound ligand/reference complex. Repeated ligand-template pairs represent different reference definitions. Composite ratios indicate shape:IFP weighting. IFD, induced-fit docking; IFP, interaction fingerprint; MD, molecular dynamics.

**Supplementary Table. S2 Static descriptors and normalized MD-quality scores for selected poses in group-matched retrospective systems**

| Ligand-template system | Group | Docking scenario | Pose-selection criterion | Static shape similarity | Static IFP score | Normalized MD-quality score (%) |
|---|---|---|---|---|---|---|
| Astemizole–8ZYO | I | Self-docking | Highest static shape similarity | 0.860 | 0.818 | 97.4 |
| Pimozide–8ZYQ | I | Self-docking | Highest static shape similarity | 0.821 | 0.914 | 95.6 |
| Pimozide–8ZYO | I | Ligand-referenced cross-docking | Highest static shape similarity | 0.737 | 0.712 | 86.6 |
| Astemizole–8ZYQ | I | Ligand-referenced cross-docking | Highest static shape similarity | 0.806 | 0.671 | 94.5 |
| Pimozide–8ZYO | I | Template-referenced cross-docking | Highest static shape similarity | 0.667 | 0.874 | 88.9 |
| Astemizole–8ZYQ | I | Template-referenced cross-docking | Highest static shape similarity | 0.680 | 0.949 | 98.0 |
| E4031–8ZYP | II | Self-docking | 30:70 IFP-favored composite | 0.772 | 0.648 | 79.4 |

Static shape similarity is the shape Tanimoto coefficient; the static IFP score measures agreement with the reference residue interaction fingerprint. Values refer to the selected pose and the reference specific to each docking scenario. Normalized MD-quality scores are expressed as a percentage of the corresponding reference MD-quality score.

The 30:70 IFP-favored composite is a provisional Group II criterion. The dual MD-quality criterion requires an absolute score ≥75 and a normalized score ≥85%. IFP, interaction fingerprint; MD, molecular dynamics.

**Supplementary Table. S3 Static similarity scores for prospective ligands across ligand-bound hERG receptor templates**

| Ligand | Ligand group | Receptor template | Template group | Group-match status | Pose-selection criterion | Static shape similarity | Static IFP score |
|---|---|---|---|---|---|---|---|
| Dofetilide | II | 8ZYP | II | Group-matched | 30:70 IFP-favored composite | 0.554 | 0.961 |
| Dofetilide | II | 8ZYO | I | Group-mismatched | 30:70 IFP-favored composite | 0.565 | 0.959 |
| Dofetilide | II | 8ZYQ | I | Group-mismatched | 30:70 IFP-favored composite | 0.535 | 0.955 |
| Ibutilide | II | 8ZYP | II | Group-matched | 30:70 IFP-favored composite | 0.435 | 0.948 |
| Ibutilide | II | 8ZYO | I | Group-mismatched | 30:70 IFP-favored composite | 0.642 | 0.951 |
| Ibutilide | II | 8ZYQ | I | Group-mismatched | 30:70 IFP-favored composite | 0.585 | 0.955 |
| Terfenadine | I | 8ZYO | I | Group-matched | Highest static shape similarity | 0.660 | 0.784 |
| Terfenadine | I | 8ZYQ | I | Group-matched | Highest static shape similarity | 0.707 | 0.938 |
| Terfenadine | I | 8ZYP | II | Group-mismatched | Highest static shape similarity | 0.545 | 0.720 |

Static shape similarity and static IFP scores refer to the selected pose and the bound ligand/reference complex of each receptor template. References differ between templates; these values should not be interpreted as directly comparable cross-template affinity or template-ranking scores.

Group-match status follows the study-defined chemical groups: 8ZYO and 8ZYQ are Group I receptor environments; 8ZYP is Group II. Terfenadine is group-matched with both 8ZYO and 8ZYQ. The 30:70 ratio indicates shape:IFP weighting and remains provisional for Group II. IFP, interaction fingerprint.
